# Type of conjugative pili governs transfer efficiency in liquid and affects interpretation of transfer assays

**DOI:** 10.64898/2026.04.01.715510

**Authors:** Nivethanaa Pulavan, Anja Nenninger, Joseph Mbuli, Julianna Poklembova, Tatiana Dimitriu

## Abstract

Plasmid conjugation is central to plasmid maintenance and spread among bacteria. Conjugation assays in liquid or on solid media are commonly used to quantify plasmid conjugation rates. Plasmids with short, rigid conjugative pili are thought to conjugate more efficiently on surfaces, whereas plasmids encoding long, flexible pili can conjugate efficiently in liquid medium. However, this pattern has not been tested systematically. Here, we perform standardised conjugation assays on a collection of 13 conjugative plasmids belonging to families that play a key role in AMR transmission and encode different conjugative pili types. We confirm that only the plasmids encoding long flexible pili conjugate efficiently in liquid. Furthermore, most transconjugants that arise from liquid assays involving plasmids with short, rigid pili can be attributed to transfer happening after the assay itself, on the surface of selective plates. This effect is amplified when using auxotrophic rather than antibiotic resistance markers, and impacts measures of transfer and defence efficiency. Finally, most of the tested plasmids with short pili had very high conjugation rates on surfaces, suggesting their transfer is mostly limited by physical constraints.

## Introduction

Horizontal transmission via conjugation is a crucial property of conjugative plasmids, responsible for the spread of plasmid-encoded traits within and among bacterial species, sometimes across large phylogenetic distances. Conjugative plasmids commonly carry accessory genes that benefit host bacteria (1,2). In particular, a few major plasmid families are now the primary vectors of antimicrobial resistance (AMR) and play a key role in the spread of AMR to pathogenic bacteria, and across environments (3–5). Models and experiments have shown that AMR plasmids can persist and spread in a population of bacteria in the absence of antibiotic selection, due to conjugation compensating for plasmid cost and loss (6,7).

Multiple genetic and environmental factors are known to impact conjugation rates (8). Plasmid, donor and recipient genotypes all contribute to variation in conjugation rates (9). For given plasmid and host genotypes, experimental conditions also strongly impact conjugation rates in a way that is dependent on plasmid properties. Early studies of a few model plasmids determined that plasmids encoding short, rigid conjugative pili transfer much more efficiently on surfaces, whereas for plasmids with flexible pili, conjugation efficiencies in liquid and on surfaces are more comparable (10,11). Studies across prokaryotic genomes have since shown that mating pair formation (MPF) systems of conjugative elements can be classified into eight classes of type IV secretion systems (T4SS), with three of them covering most conjugative plasmids in Proteobacteria (12–14). MPF_T_ encode short, rigid pili and transfer efficiently on solid surfaces; MPF_F_ encode long, flexible pili that also allow mating pair stabilisation and efficient transfer in liquid (15). Finally, MPF_I_ encode short, rigid pili similar to MPF_T_ systems, but are usually carried on plasmids also encoding a thin pilus homologous to type IV pili (16,17). Expression of these thin pili either in the donor or the recipient can rescue transfer of MPF_T_-encoding pili in liquid by stabilising mating pairs (18).

Transfer in liquid or on solid surfaces has been compared quantitatively for a few model plasmids, almost exclusively IncF plasmids for MPF_F_ and IncP for MPF_T_ types (19,20). Meta-analyses suggest that ‘media type’ (including liquid, plate, filter and biofilm assays) has a strong impact on transfer efficiency (9), but this pattern has not been tested with quantitative comparison across plasmid types. Moreover, conjugation assays also vary in the method used to quantify the cell densities of plasmid donors, recipients and transconjugants (recipients which received the plasmid) at the end of the mating assay (Fig. 1). Plasmid carriage in donors and transconjugants is usually identified by plating onto antibiotics to which plasmid-encoded AMR genes confer resistance. To distinguish the donor from the recipient strain, two main methods are used, both of which counter-select donors. Recipients can first be marked with an antibiotic resistance marker, usually selecting for spontaneous resistance to rifampicin or to nalidixic acid (Nal). Nal resistance is more often used, as it can also inhibit conjugation from sensitive donors (21) although that depends on the plasmid considered (22). Alternatively, a more recent method uses engineered *dapA* knockout donors, which are auxotrophic for diaminopimelate (DAP) and unable to grow even in rich media without DAP supplementation (23,24). This method is convenient when screening large numbers of recipients, as one engineered donor can be used in combination with native recipients. However, donors will not die on transconjugant selective plates and might engage in further transfer events. A recent study demonstrated that for plasmid RP4, transfer on transconjugant-selective plates can happen at non-negligible rates (in this case, using Nal resistance for selecting recipients), because selection against donors and recipients is not effective or rapid enough (25). In turn, this strongly inflates measures of RP4 transfer efficiency from assays performed in liquid conditions, where RP4 transfers poorly. We expect that the same issue might affect other plasmids with preferential transfer on solid surfaces and be compounded when using DAP auxotrophy to counter-select donors.

**Figure 1:**
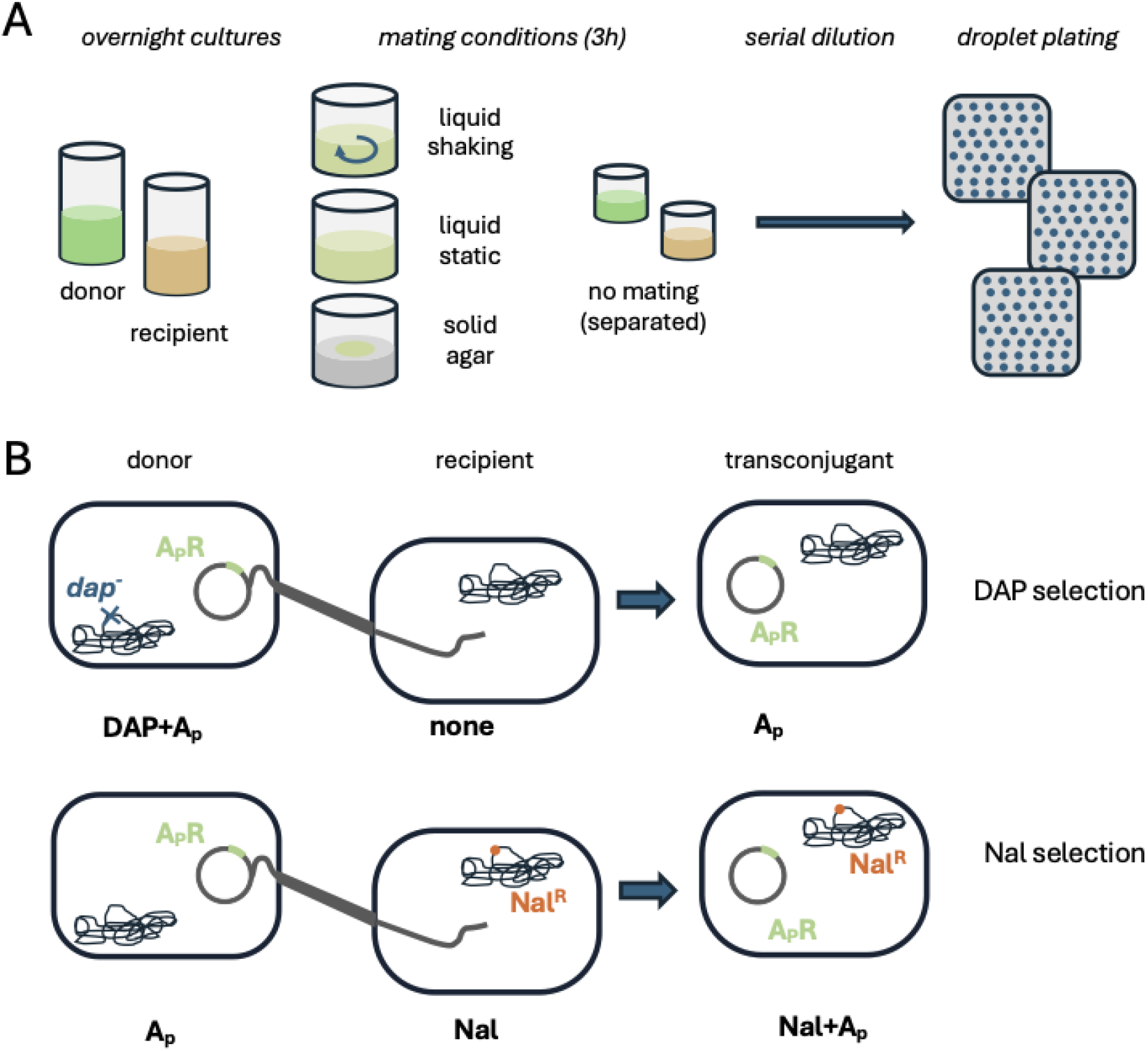
design of conjugation assays and conditions tested. **A: mating conditions**. Donors and recipients were mixed for 3h in liquid with or without shaking or on agar; a control treatment without mating (“no mating” treatment) was performed in parallel by incubating donors and recipients separately. **B: phenotypic markers used to distinguish cell types**. Plasmids were identified by one of the resistance markers they carry (see Table 1). Recipients were distinguished from donors by their ability to grow in the absence of DAP (DAP method) or in the presence of Nal (Nal method).

Here, we compare conjugation efficiency in liquid and on solid surfaces for 13 conjugative plasmids belonging to major families of plasmids that spread AMR within Enterobacteriaceae and include the three most common types of conjugative pili. We also compare alternative methods for distinguishing donors from recipients.

**Table 1:** conjugative plasmids used, their relevant properties and antibiotics used for selection. Relaxase and MPF type were annotated using MOB-suite. ^*^ indicates that the *pil* operon is also present.

| Plasmid | Inc group | relaxase | MPF type | Selection |
| --- | --- | --- | --- | --- |
| R1 | F | MOBF | MPF_F | Kan |
| pRK100 | F | MOBF | MPF_F | Tc |
| pKAZ3 | C | MOBH | MPF_F | Tc |
| pCT | K | MOBP | MPF_I* | Amp |
| pESBL283 | I | MOBP | MPF_I* | Strep |
| pOXA-48 | L | MOBP | MPF_I | Amp |
| R6K | X | MOBP | MPF_T | Strep |
| pCF12 | X | MOBP | MPF_T | Strep |
| pCU1 | N | MOBF | MPF_T | Strep |
| RIP113 | N | MOBF | MPF_T | Tc |
| pB10 | P | MOBP | MPF_T | Tc |
| RP4 | P | MOBP | MPF_T | Tc |
| R388 | W | MOBF | MPF_T | Tm |

## Materials and methods

### Bacterial strains, plasmids and growth conditions

Plasmid donors were MG1655 or its variant MG1655 Δ*dapA*::ErmR described in (26); plasmid recipients were MG1655, MG1655 Nal^R^ (a spontaneous mutant resistant to nalidixic acid), MG1655 Nal^R^ pEcoRV (carrying the EcoRV restriction-modification system, (26)) and 4 natural isolates of *E. coli* described in (27) and marked with rifampicin in (28). The 13 conjugative plasmids studied are listed in Table 1.

Bacterial strains were grown in LB (Miller’s LB Broth) growth medium, with 150 rpm shaking at 37°C. For solid medium 1.5% agar was added. Antibiotics were added at the following concentrations: ampicillin (Amp, 100 μg/mL), chloramphenicol (Chl, 20 μg/mL), kanamycin (Kan, 50 μg/mL), nalidixic acid (Nal, 30 μg/mL), streptomycin (Strep, 50 μg/mL), tetracycline (Tc, 10 μg/mL), and trimethoprim (Tm, 10 μg/mL). 2,6-diaminopimelic acid (DAP) was added to Δ*dapA* cultures at 0.3 mM for overnight cultures and plates, and at 0.15 mM for conjugation assays.

### Conjugation assays

For conjugation assays, donor and recipient strains were first grown in the absence of antibiotics, except MG1655 Nal^R^ pEcoRV which was grown with Amp to maintain pEcoRV. Conjugation assays were performed in 24-well plates. For both liquid mating treatments, 75 μL each of the donor and recipient cultures were added into 900 μL of pre-warmed LB. For the solid treatment, 500 μL each of the donor and recipient strains were mixed and centrifuged at 3500rpm for 2 minutes. The pellet was then resuspended in 1ml fresh LB broth and 15 μL of this resuspension was spotted onto pre-warmed LB agar. Following (29), this method maximises contact between donor and recipient cells and minimises the complexity of growth on surfaces. If initial densities on agar are too low, transfer becomes largely dependent on the limited contact between separated donor and recipient microcolonies (19,30,31).

In the case of Δ*dapA* donors, DAP was added at final concentration of 0.15 mM. All mating cultures were incubated for 3 hours at 37°C, under static conditions for liquid static and agar treatments and at 150 rpm shaking for the liquid shaken treatment (Fig. 1A). After 3 hours, conjugation treatments on agar were first resuspended in 1 mL PBS by pipetting, then all treatments were serially diluted in PBS. Controls in which donor and recipient strains were grown alone for 3 hours were included under all three conditions, and after 3 hours, these donors and recipients were mixed then immediately serially diluted, in order to evaluate the amount of conjugation happening on selective plates (no mating treatment).

Two donor and recipient combinations using different selection methods were compared (Fig. 1B). In the ‘dap’ method, donors were unable to grow in the absence of DAP supplementation; in the ‘nal’ method, donors were sensitive to nalidixic acid. Appropriate dilutions were plated on selective medium using 10 μL droplets: for experiments using Δ*dapA* donors and *dapA*^*+*^ recipients, dilutions were plated on LB-agar + A_p_ + DAP, LB-agar and LB-agar + A_p_ to estimate densities of donors, recipients and transconjugants respectively, where A_p_ stands for the antibiotic used to select for a given conjugative plasmid (see Table 1). For experiments using Nal-sensitive donors and Nal^R^ recipients, selective plates were LB-agar + A_p_, LB-agar + Nal and LB-agar + Nal + A_p_. By default, dilutions 10^−3^ to 10^−6^ were plated for donors and recipients, and 10^0^ (undiluted) to 10^−6^ for transconjugants, with the 10^−1^ dilution repeated twice to increase the limit of detection (the undiluted droplets frequently were uncountable due to poor selection at high cell density). For plasmid R6K, which has low transfer efficiency, 100 μL of the undiluted conjugation was also plated separately and the incubation time for the assay was extended to 4 hours at 37°C. All experiments were run in six replicates

### Growth rate measurements

We measured the exponential growth rate of cells in conjugation assay settings in two ways. First, 6 replicates of the MG1655 Nal^R^ recipient were grown under the conditions of the liquid shaking mating treatment, and OD600 readings were taken in a NanoPhotometer NP80 (GeneFlow) reader every 30 minutes (Fig. S1A). Exponential growth rate was calculated manually during exponential growth from the 1h and 4h readings, yielding an average growth rate of r = 1.06 h^-1^.

To screen for differences in growth rate across donor strains, we measured exponential growth rate in a plate reader. Overnight cultures were diluted 100-fold into 200 µL LB in 96-well microplates and covered with 40 µL mineral oil. Optical density at 600 nm was measured at 3-minute intervals in a Multiskan SkyHigh plate reader at 37 °C for 16h with shaking on. Maximal growth rate was computed using the R package growthrates between 1h and 5h post-inoculation, with the h parameter set at 8. Some plasmids imposed a cost on the host, especially in the *dapA* strain (Fig. S1B). However, as differences across strains were not large and rarely reproducible across two independent experiments (Fig. S1B), we used r = 1 h^-1^ as an estimate of growth rate across experiments for calculating transfer rates below. Small differences in donor growth rates should lead to limited impacts on calculated transfer rates (32).

### Data analysis

All statistical analyses were performed with R version 4.3.2 (33). Packages reshape (34) and cowplot (35) were used for data processing and plotting. To calculate transfer rates, we first excluded rare outliers for which the density of donor or recipient cells was lower than 10^4^ cells/μL. Of note, transfer assays with pOXA-48 plasmid in the *dapA* donor had to be repeated, as several independent attempts yielded no viable counts for donors on LB + Amp + DAP (despite transconjugant colonies being present). When no transconjugant colony was present, a threshold transfer efficiency was calculated by assuming that 0.5 transconjugant colony was detected. Transfer rates were calculated using the endpoint method (36). For significance testing, all transfer and cell density measures were first log-transformed.

## Results

We performed conjugation assays for 13 conjugative plasmids, each with two selection methods (DAP or Nal), and four mating conditions (no mating treatment, liquid + 150 rpm shaking, liquid static, solid). We first analysed the results obtained using the Nal method (Figure 2). Across plasmids, we observed a large effect of mating conditions on measured transfer rates, and this effect interacts with plasmid identity (Fig. 2A; log_10_ transfer rate ∼ treatment × plasmid, treatment effect F_3,258_ = 513, *p* < 2 × 10^−16^, plasmid effect F_12,258_ = 58.2, *p* < 2 × 10^−16^, interaction effect F_36,258_ = 22.8, *p* < 2 × 10^−16^). Overall, the lowest transfer rate was observed for the no mating treatment, then increasing transfer for liquid static, liquid shaking and finally solid (TukeyHSD, all *p* < 0.002). However, this pattern varied across plasmids (Fig. 2A and B).

**Figure 2:**
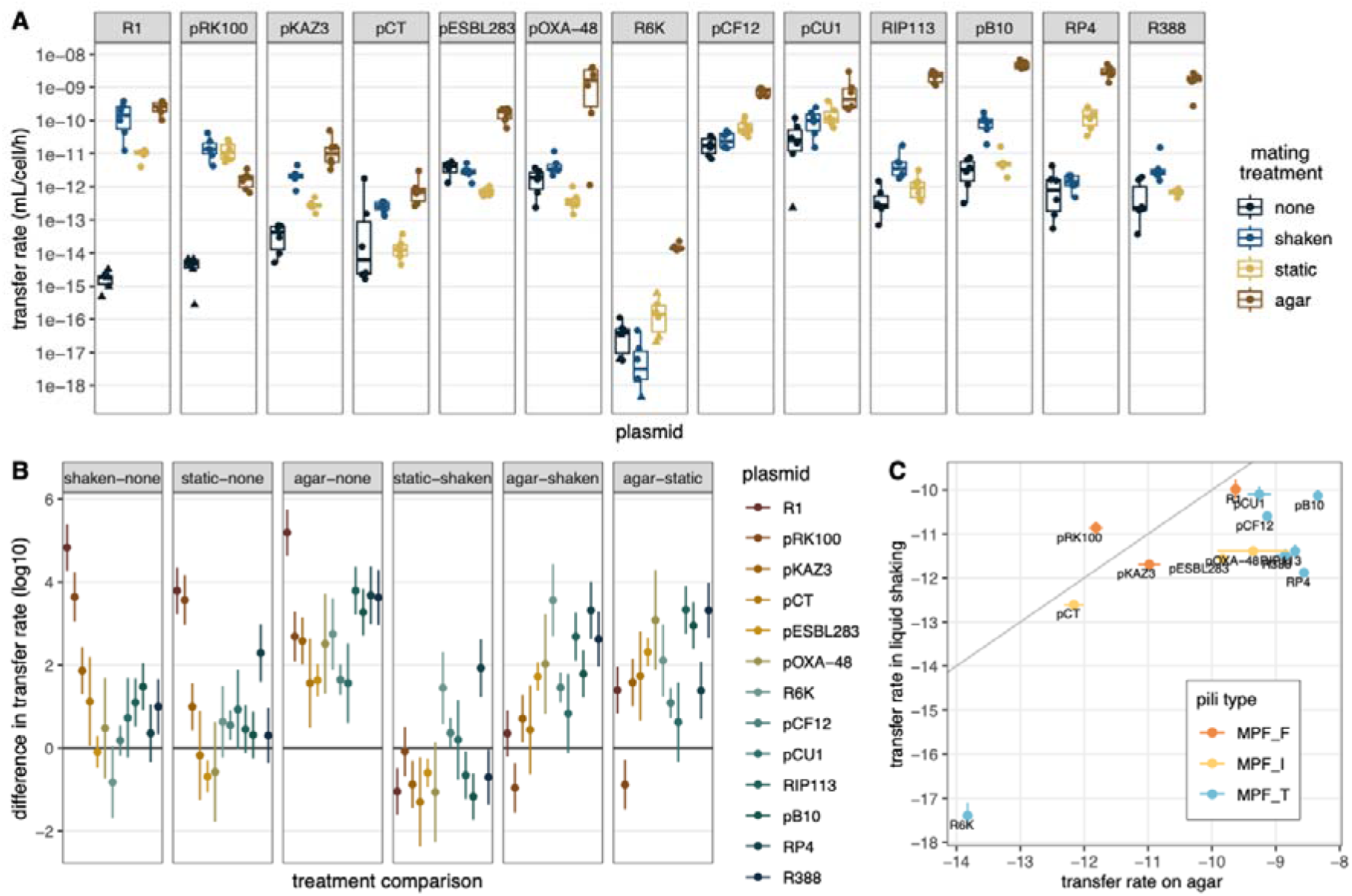
Impact of mating conditions on plasmid transfer rates using Nal selection. A shows plasmid transfer rates in the 4 different mating conditions for 13 conjugative plasmids. The centre line of the boxplots shows the median, boxes show the first and third quartiles, and whiskers represent 1.5 times the interquartile range; individual data points are shown as dots (n=6). Triangles indicate replicates for which no transconjugant CFUs were observed and transfer rates were computed using a threshold transconjugant density. B shows the summary of pairwise comparisons between treatments for each plasmid (TukeyHSD tests, dots show the difference in means, lines show 95% confidence interval). C shows transfer rates in liquid shaking conditions as a function of transfer rates on solid surfaces, with dots and lines showing respectively the average and standard error, respectively. The grey line indicates equal transfer rates in both conditions, colour shows annotated MPF type.

Mating in liquid yielded transfer rates higher than no mating treatments only for 7 of 13 plasmids in shaking, and 5 of 13 in static conditions. In contrast, all 13 plasmids transferred at higher rates on solid compared to no mating treatments. Moreover, the majority of plasmids had higher transfer rates on solid agar compared to liquid conditions. This is consistent with the hypothesis that conjugation in no mating treatments happens after plating on agar and plasmids with low transfer rates in liquid will thus not differ significantly from the no mating treatment. Across plasmids, preferential transfer on agar was found mostly for plasmids encoding MPF_T_ type pili (Fig. 2C). In contrast, all plasmids encoding MPF_F_ type pili transferred at similar rates in liquid and on agar. MPF_I_ -type plasmids were less consistent, with pCT transferring equally in both conditions, whereas pESBL283 and pOXA-48 transferred more in liquid.

Conjugation assays using DAP for selection led to globally similar patterns (Fig. S1), except that static and shaking liquid mating treatments were not significantly different across plasmids. There were some relevant differences for individual plasmids. First, pCF12 plasmid transferred at noticeably higher rates across mating conditions (this effect was repeatable in independent experiments). There were also fewer significant differences between treatments (Fig. S1B). Most notably, 18 comparisons to the no mating treatment (of 39 possible comparisons) were not significantly different, up from 13 comparisons with Nal selection. Liquid shaking transfer rates were only significantly increased compared to no mating conditions for the three plasmids encoding MPF_F_ pili. This is all consistent with counter-selection of mating partners on selective plates being less effective with DAP selection. We next zoom in on results from the ‘no mating’ treatment to analyse this in more detail.

Across plasmids, DAP selection led to higher transconjugant densities in the no mating treatment (Fig. 3A; log_10_ transconjugant density ∼ selection method × plasmid, selection effect F_1,127_ = 46.6, *p* < 4 × 10^−10^, plasmid effect F_12,127_ = 18.4, *p* < 2 × 10^−16^, interaction effect F_12,127_ = 18.4, *p* < 2 × 10^−16^). The two exceptions were pKAZ3 and pCF12 (but the latter has genuinely higher transfer rates across treatments). As plasmids also vary in the resistance markers we use for selection, which are likely to act differently on recipients, we next wondered whether these contribute to variation in the amount of conjugation happening on selective plates. To test this while controlling for plasmid intrinsic transfer rate, we performed ‘no mating’ assays, using different antibiotics for selection for two plasmids carrying multiple resistance markers. R1 produces MPF_F_ pili and transconjugants can be selected using Amp, Chl, Kan or Strep, and the MPF_T_ pili producing RP4 can be selected using Amp, Kan or Tc. For both plasmids, we observed a strong impact of the method used for counter-selecting donors (Fig. 3B). Nal selection was always at least as effective, and, in most cases, more effective than DAP selection. For both plasmids, Amp was the least effective antibiotic, and the measured transfer rates were highest for RP4 using DAP and Amp selection. Apart from Amp, bactericidal antibiotics (kanamycin and streptomycin) were more effective than bacteriostatic antibiotics (chloramphenicol and tetracycline). Differences between methods led to almost 5 orders of magnitude variation in recorded transfer rates for RP4, and 3 orders of magnitude for R1. Thus, the choice of markers can be critical to limit artefacts of transfer on selective plates for all plasmids, but especially ones with preferential transfer on surfaces.

**Figure 3:**
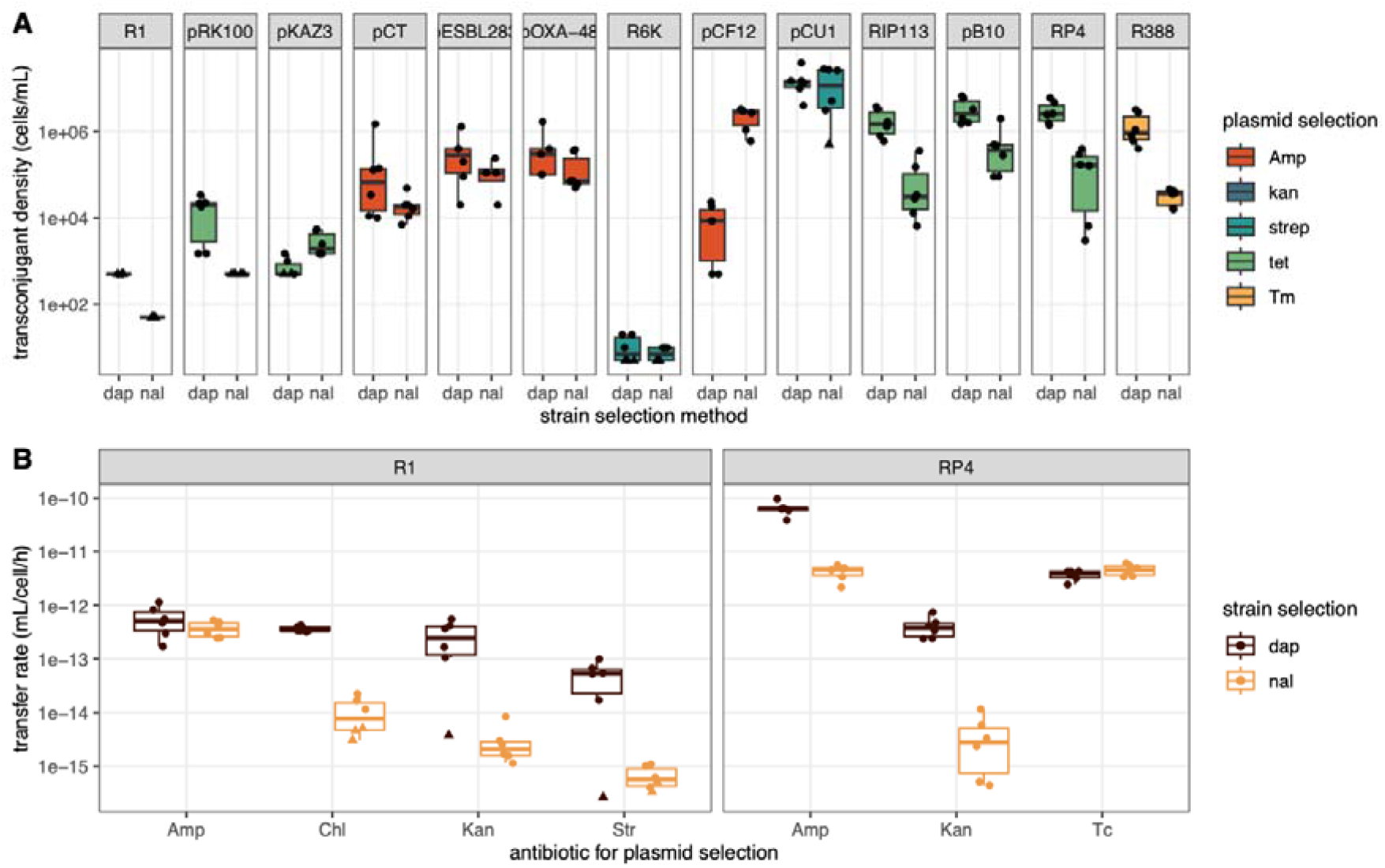
Effect of strain and plasmid selection methods on measured transfer rates in the absence of mating before plating. A shows transconjugant density for the ‘no mating’ treatment across 13 conjugative plasmids, for both the DAP and Nal methods. In B, an independent ‘no mating’ assay was performed for plasmids R1 and RP4, and selective plating used one of four (R1) or three (RP4) different plasmid selective markers. The centre line of the boxplots shows the median, boxes show the first and third quartiles, and whiskers represent 1.5 times the interquartile range; individual data points are shown as dots (n=6). Triangles indicate replicates for which no transconjugant CFUs were observed and transfer rates were computed using a threshold transconjugant density.

Finally, we revisited some previous results on RP4 transfer efficiency towards natural isolates (28), and towards recipients carrying restriction-modification systems (26), which were obtained in liquid shaking conditions and using DAP selection. Comparing liquid and agar mating, we observed a strong effect of mating conditions on transfer rates, and this effect interacted significantly with the identity of recipient strains (Fig. 4, log_10_ transfer rate ∼ recipient × treatment, recipient effect F_5,34_= 51.9, *p* < 6 × 10^−15^, treatment effect F_1,34_= 242, *p* < 2 × 10^−16^, interaction effect F_5,34_= 17.1, *p* < 2 × 10^−8^). This impacts not only absolute measures of transfer, but also conclusions about relative transfer or defence efficiency. For instance, carrying the EcoRV RM system was associated with a 10000-fold decrease in measured transfer in liquid, and only a 10-fold decrease on agar. A similar behaviour was present for natural isolates D7.8 and R3.1. This is likely due to saturation of transfer in agar mating: effectively all MG1655 and R3.1 recipients received the plasmid, limiting the detection of stronger differences. In contrast, oc1.5 and R1.9 isolates still received few plasmids in agar treatments, suggesting they present a more robust barrier to transfer, and lead to a higher relative rate of transfer compared to MG1655. Thus, conditions of conjugation assays have a direct impact on absolute and relative variation in transfer across strains.

**Figure 4:**
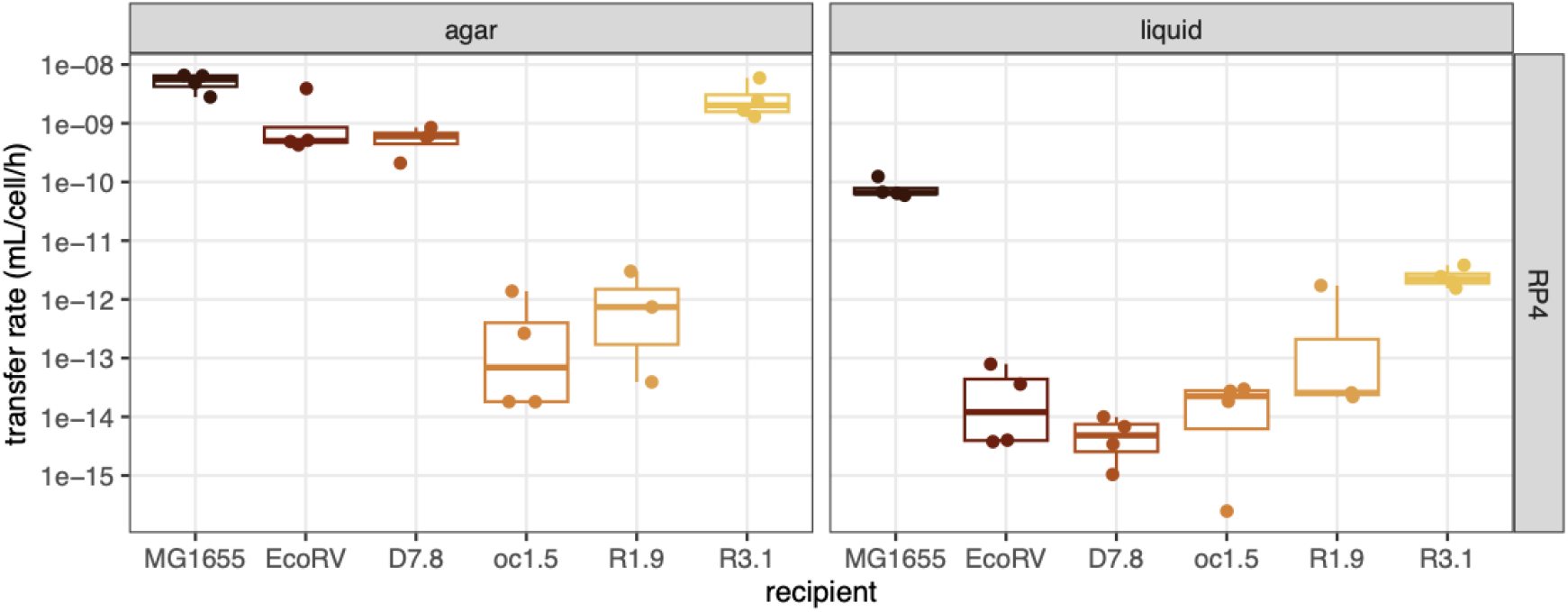
Mating condition impact measures of recipient ability and defence efficiency. Absolute transfer rates from MG Δ*dapA* RP4 towards different *E. coli* recipients are shown for mating assays on agar or in liquid. The centre line of the boxplots shows the median, boxes show the first and third quartile, and whiskers represent 1.5 times the interquartile range; individual data points are shown as dots (n=4).

## Discussion

Here, we investigated the impact of mating conditions on plasmid conjugation. Overall, our results align with expectations about the role of different pili types in liquid or solid mating: plasmids with long, flexible MPF_F_ pili transferred well in liquid, whereas plasmids with short, rigid MPF_T_ pili transferred well only on agar. MPF_I_ plasmids were more variable: pOXA-48 preferential transfer on agar is consistent with MPF_I_ short, rigid pili behaving similarly to MPF_T_ pili. pCT and pESBL283 additionally produce thin pili encoded by the *pil* locus, which should facilitate liquid mating, yet only pCT transferred efficiently in liquid conditions. This might be due to the presence within the *pil* locus of a locus prone to frequent rearrangements called a shufflon, and which has different versions that determine recipient specificity in liquid (37,38). The shufflon configuration in pESBL283 might not be optimal for transfer towards MG1655, making this plasmid behave similarly to a plasmid that does not express long flexible pili.

On agar, most MPF_T_ plasmids (apart from IncX plasmids) displayed extremely high transfer rates around 10^−9^ mL/cell/h. These are the maximal rates achievable in these assays, as they correspond to most recipient cells acquiring the plasmid. Thus, shorter assays would be needed to estimate transfer rates more accurately, and real transfer rates are likely to be even higher. These high transfer rates suggest that most plasmids with MPF_T_ pili are highly transfer proficient. Indeed, IncP plasmids do not display on/off expression of transfer genes but instead all cells appear to be able to transfer plasmids due to low but tightly regulated, and constant levels of transfer gene expression (39). IncX plasmids might diverge from this pattern with an on/off regulation more similar to F-like plasmids (40).

In liquid conditions, transfer efficiency is limited by the rates of donor-recipient attachment, mating pair formation and stabilisation (30,41). For plasmids with MPF_T_ pili, rates measured after mating in liquid were much lower than after agar mating, and often not significantly different from the no mating treatment. Thus, most observed transfer actually arises from transfer happening on the selective plates. This suggests that for plasmids encoding MPF_T_ pili, attachment and transfer are strongly limited, if not impossible, in liquid environments. In contrast, for MPF_F_-encoding plasmids, we measured similar rates of transfer in liquid and on agar, confirming that MPF_F_ pili are extremely effective at mediating attachment even with high levels of shear (15), and attachment is not the limiting factor. MPF_F_ plasmids instead tend to repress transfer gene expression in the majority of cells, as described in detail for F-like plasmids (42) and also characterised for IncC plasmids (43). Plasmids with mutations inactivating the repressor (44) or increased expression of the transfer operon due to increased copy number (45) invade populations of recipients in liquid much faster (45,46), consistent with the idea that repression is the main limiting factor for their transfer.

Finally, the effect of shaking was not strong and was not consistent across plasmids. Our results suggest that using static conditions to prevent pili from breaking, a common assumption of some transfer assays (11), is unlikely to be crucial to the success of transfer assays. Detailed effects of shaking are likely to depend on the exact amount of shear generated by mixing (47), its differential effects on attachment and detachment (41), and the initial density of cells (48).

We further show that preferential transfer on agar affects the results of transfer assays done in liquid, by leading to high background levels of transconjugants arising after plating and not during the assay. Our results generalise previous work using plasmid RP4 (25,49) to any plasmid with preferential transfer on agar. Transfer on selective plates can be reduced by several orders of magnitude by using nalidixic acid to counter-select donors. This underlies the limitation of using DAP auxotrophy as a counter-selection method: auxotrophs will still survive on agar and actively transfer even they cannot grow to form visible colonies. Instead, Nal inhibits conjugation from some donors (22,50), and also likely acts more directly by killing the sensitive donors. Different plasmid selection antibiotics also vary in their effect on plasmid-free recipients: we observe that bactericidal antibiotics are more effective than bacteriostatic ones, as expected, as death of recipients will prevent subsequent establishment of plasmids on transconjugant plates. Ampicillin was the exception, which can be explained as B-lactamase production is cooperative and protects sensitive recipients, allowing subsequent transfer (51).

More generally, for plasmids with preferential transfer on agar, the choice of mating assays will impact conclusions that can be drawn. Agar matings should be more accurate, as any treatment applied during the mating treatment will not be sustained after plating on transconjugant plates, where further transfer will confuse the results. On the other hand, the duration of mating might need to be reduced further to prevent full invasion of the recipient by plasmids. When agar matings are not practical, Nal selection and cidal antibiotics can at least limit background transfer on agar. Simply plating transconjugants on full Petri dishes will also limit this by dilution, limiting contact between donor and recipient cells. Alternatively, to measure transfer rates accurately for a few specific clones or conditions, the Luria-Delbrück fluctuation approach can be used (52). Ultimately, to understand transfer in natural and clinical contexts, it is necessary to account for the complexity of natural environments like the gut microbiome (53). Transfer on agar and in well-mixed liquid conditions are likely to be only simplified extreme conditions, but pili type has been shown to play an important role in the gut, with the mating pair stabilisation of IncI2 plasmids being particularly effective at transfer in the gut (38,54). Understanding in which conditions various types of plasmids spread will be important both to understand AMR plasmid spread, but also for engineering plasmids that target AMR plasmids or strains (56,57).

## Author contributions

NP, AN, JM, JP, TD: Investigation, Methodology. NP, AN, TD: Writing. TD: Conceptualization, Funding acquisition, Supervision.

## Conflicts of interest

The authors declare that there are no conflicts of interest.

## Funding information

This work was funded by the Royal Society (University Research Fellowship URF\R1\231740) https://royalsociety.org/ to T.D. The funders played no role in the study, preparation of the article or decision to publish.

## Acknowledgments

We thank Andy Matthews for initial discussions about experiment design, and Allan Zuza for help with plasmid typing.

## Notes

### Competing Interest Statement

The authors have declared no competing interest.

