## Supplementary information for "Type of conjugative pili governs transfer efficiency in liquid and affects interpretation of transfer assays"

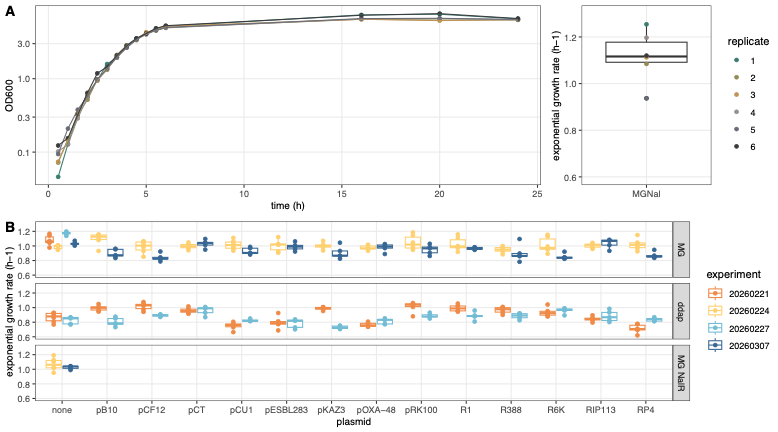
**Figure S1:** **exponential growth rate measures.** A shows growth curves of MG1655 Nal^R^ measured manually and associated exponential growth rate; B shows exponential growth rate from all plasmid donor and recipients measured in 96-well plates in a plate reader. The centre line of the boxplots shows the median, boxes show the first and third quartile, and whiskers represent 1.5 times the interquartile range; individual data points are shown as dots (n≥4).

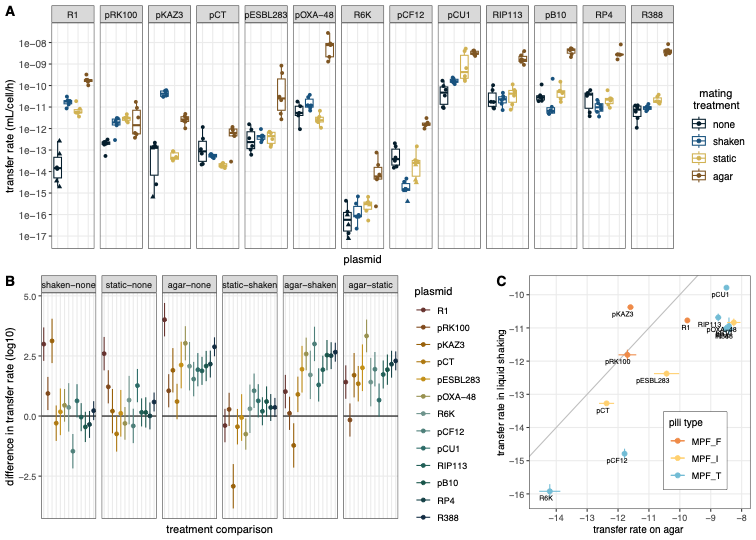

**Figure S2:** **Impact of mating conditions on plasmid transfer rates using DAP selection**. A shows plasmid transfer rates in the 4 different mating conditions for 13 conjugative plasmids. The centre line of the boxplots shows the median, boxes show the first and third quartile, and whiskers represent 1.5 times the interquartile range; individual data points are shown as dots (n=6). Triangles indicate replicates for which no transconjugant CFUs were observed and transfer rates were computed using a threshold transconjugant density. B shows the summary of pairwise comparisons between treatments for each plasmid (TukeyHSD tests between treatments for each plasmid, dots show the difference in means, lines show 95% confidence interval). C shows transfer rates in liquid shaking conditions as a function of transfer rates on solid surfaces for each plasmid, with the grey line indicating equal transfer rates in both conditions. Colour indicates annotated MPF type. Dots and line show respectively the average and standard error per plasmid.
